# EEG signatures of contextual influences on visual search with real scenes

**DOI:** 10.1101/2020.10.08.332247

**Authors:** Amir H. Meghdadi, Barry Giesbrecht, Miguel P Eckstein

**Affiliations:** Department of Psychological and Brain Sciences, University of California, Santa Barbara, Santa Barbara, CA, 93106-9660; Institute for Collaborative Biotechnologies, University of California, Santa Barbara, Santa Barbara, CA, 93106-5100; Interdepartmental Graduate Program in Dynamical Neuroscience, University of California, Santa Barbara, Santa Barbara, CA, 93106-5100

**Author notes:** **Corresponding author:** Amir H. Meghdadi, Department of Psychological and Brain Sciences, University of California, Santa Barbara, Santa Barbara, C, 93106-9660.

**Keywords:** EEG, ssVEP, visual search

## Abstract

The use of scene context is a powerful way by which biological organisms guide and facilitate visual search. Although many studies have shown enhancements of target-related electroencephalographic activity (EEG) with synthetic cues, there have been fewer studies demonstrating such enhancements during search with scene context and objects in real world scenes. Here, observers covertly searched for a target in images of real scenes while we used EEG to measure the steady state visual evoked response to objects flickering at different frequencies. The target appeared in its typical contextual location or out of context while we controlled for low-level properties of the image including target saliency against the background and retinal eccentricity. A pattern classifier using EEG activity at the relevant modulated frequencies showed target detection accuracy increased when the target was in a contextually appropriate location. A control condition for which observers searched the same images for a different target orthogonal to the contextual manipulation, resulted in no effects of scene context on classifier performance, confirming that image properties cannot explain the contextual modulations of neural activity. Pattern classifier decisions for individual images was also related to the aggregated observer behavioral decisions for individual images. Together, these findings demonstrate target-related neural responses are modulated by scene context during visual search with real world scenes and can be related to behavioral search decisions.

**Significance Statement:** Contextual relationships among objects are fundamental for humans to find objects in real world scenes. Although there is a larger literature understanding the brain mechanisms when a target appears at a location indicated by a synthetic cue such as an arrow or box, less is known about how the scene context modulates target-related neural activity. Here we show how neural activity predictive of the presence of a searched object in cluttered real scenes increases when the target object appears at a contextual location and diminishes when it appears at a place that is out of context. The results increase our understanding of how the brain processes real scenes and how context modulates object processing.

## Introduction

Humans and other animals have a remarkable ability to visually search for targets in real scenes. One important strategy utilized by many species is to rely on statistical properties and other elements/objects of scenes predictive of the target location to guide search (Bushnell and Rice 1999; Castelhano and Heaven 2010; Chun and Jiang 1998; Eckstein 2013; Wasserman et al 2014, Wolfe et al 2011). Thus, when a hard to see target appears spatially close to a highly visible cue (e.g., an associated object, or visual feature/s), the target is often detected faster or more accurately than when the cue is not proximal, not predictive, or absent altogether. In humans, these contextual benefits are observed when the predictive value of the spatial cue is explicitly provided to the observer (Carrasco 2011; Eckstein et al 2004; Luck 1994; Posner 1980) when it is learned (Droll et al 2009), and/or when the spatial location information is provided by the configuration of distractors in a search array (Chun 1998; Giesbrecht 2013).

When humans search real scenes, global statistical properties (Torralba, 2006; Wolfe, 2011), objects that often co-occur with the searched target (Castelhano, 2011; Eckstein, 2006; Mack, 2011; Vo, 2012; Wolfe, 2011), and the configuration multiple objects which jointly specify a likely target location (Koehler and Eckstein, 2017), all guide eye movements and facilitate visual search.

While a number of studies have investigated the neural correlates associated with context during visual search of synthetic displays (e.g., Johnson et al. 2007; Giesbrecht et al. 2013), understanding of the neural mechanisms of contextual effects in search of natural scenes is more limited. Most relevant for the present work are the handful of studies utilizing EEG while observers search for targets in real scenes. Gerson et al (2006) used EEG and pattern classifiers to identify rapidly presented images that contained a person and achieved an accuracy of 92% with a 50 ms time window of neural data. A recent study found a significant difference in the P300 component of event-related potentials (ERPs) when observers moved their eyes toward targets vs. distractors in natural scenes (Brouwer 2013; Devillez 2015). Given the important role of scene context for visual search, it is likely that the EEG signals might carry information about the contextual locations of visual targets. Previous studies have evaluated how semantic consistency of a target with the scene or its spatial location modulates ERPs (Demiral et al. 2012; Kutas and Hillyard 1980; Vo and Wolfe 2011) but in these studies, observers did not engage in a search task making it difficult to isolate EEG components directly reflect the detection of the searched target from violations of expectations.

Here we investigated the influence of scene context effects on the accuracy of EEG-based target detection while observers searched for a target in a cluttered real scene. To maximize the target-related EEG signals relative to background noise, we flickered non-target objects in real scenes (Figure 1). These objects were flickered at specific frequencies to induce stimulus-related oscillations in the EEG data, commonly known as the steady-state visually evoked potential (Reagan 1966). The location of the target object was manipulated such that in most trials the target (computer mouse) was located at an in-context (i.e., expected) location while in a small subset of trials the target object was placed out of context (unexpected). The target itself was not modulated temporally. We controlled for various low level variables across conditions including target saliency with respect to the neighboring background, the average retinal eccentricity of the target, and the size of the target. We utilized a simple pattern classifier (linear discriminant analysis, LDA) and evaluated the time course of target detection accuracy from EEG power at the stimulus flicker frequency as a function of whether the target was at the contextual location or not. To discount the possibility that differences across contextual conditions might arise due to the physical aspects of the images, we included a control condition in which observers viewed the same images but searched for a different object (stapler).

**Fig. 1.**
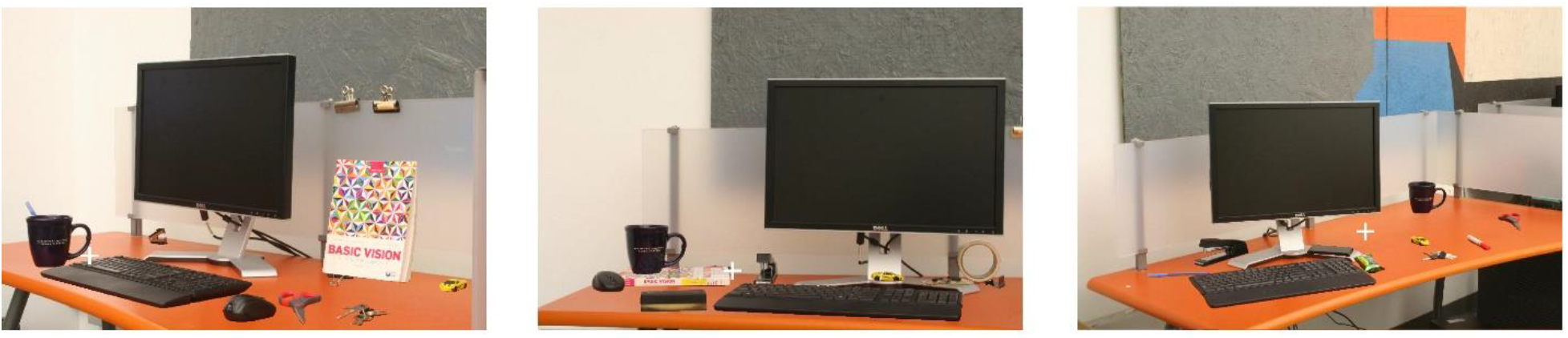
Example of an image with the target (mouse) present in context next to keyboard (left image), the target out of context at an irrelevant location (middle image) and target absent (right image). Fixation cross is always located halfway between imaginary bounding boxes encompassing the flickering objects (between keyboard and cup)

## Methods

### Participants

Fourteen subjects participated in this study (9 females and 5 males), all university students (18-24 years old) and received course credit for their participation. All participants had normal or corrected to normal vision. The data acquisition was successful with thirteen of the fourteen subjects. For one subject the EEG cap was an incorrect size leading to low-quality EEG data. The data from this subject was excluded from further data analysis. All procedures were approved by the UC Santa Barbara Human Subjects Committee.

### Stimuli

Images were generated by manually taking photos of a computer desk from multiple viewing angles and distances (24 unique configurations), with and without 16 objects randomly placed on the desk. Objects were manually segmented from the background using digital image editing tools. Finally, a set of 96 images was generated by adding a random selection of the objects to the background images. All images contained a keyboard, a cup, a monitor and a random number (between 8 to 11) of distracting objects.

On each trial, the image was redrawn at the appropriate screen refresh time (60 Hz) where the cup and keyboard were added/not added to the background and flickered with a tagged frequency of either 12.1648 Hz or 14.1923 Hz, with the purpose of eliciting a steady-state visually evoked potential (SSVEP) at each of those frequencies. Eliciting reliable SSVEPs is particularly challenging in this experiment because a) our flickering objects are natural images rather than distinct high contrast patterns (such as checkerboards) normally used in SSVEP research and b) the power of SSVEPs changes as a function of eccentricity (Regan, 1966; Ding et al., 2006; Lin et al., 2012). Therefore, we resized our displayed images such that the eccentricity of tagged objects with respect to the fixation did not exceed 5 degrees of the visual field. This number was chosen based on unpublished preliminary testing to determine the detectability of SSVEPs as a function of eccentricity. Displayed images were 652×434 pixels in size on a CRT display with a resolution of 1280 by 1024, placed at 105 cm distance from the subjects’ eyes. At this distance, the images subtended a 10° × 6.6° visual angle.

### Experimental protocol

All participants were naive to the purpose of the study. Participants sat inside a dark electromagnetically shielded booth (ETS Lindgren). Participants gave written informed consent and their participation was voluntary.

The task was to search for a target in a real scene. The experiment consisted of 576 trials divided into 6 blocks, 96 trials each. Prior to the main experiment, they were shown a demo version of the experiment to practice, which was the same as the main experiment. On each trial, a stimulus image was presented to the subject for 3 seconds. Fig. **2** shows the task flow. Before each trial, subjects were instructed to fixate on a fixation cross and press a key on the keyboard to indicate they are ready for the trial. Each trial was then started at a random time ranging from 500 and 750 ms after the key-press and lasted for 3 seconds. The subject’s task was to search for the target object while maintaining fixation on the fixation cross. After each trial, the image disappeared, and the subjects were asked to press the corresponding keys to indicate if the target was “absent” or “present” in the scene. The keys used for present and absent responses (F and J on a standard keyboard) were counterbalanced between subjects. The fixation cross was always located midway between the flickering objects (cup and keyboard). Subjects were instructed not to do anticipatory responses. The midway location was chosen to minimize the difference between the eccentricities of flickering objects with respect to the fixation point.

**Fig. 2.**
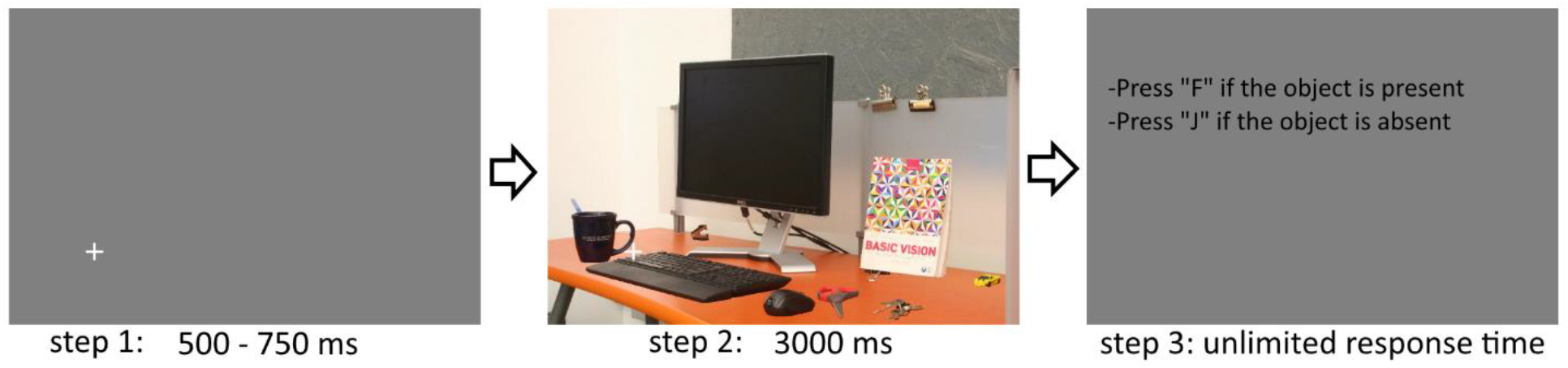
Experimental procedure for each trial: subjects fixate on a fixation cross on a gray background (step 1), the trial starts between 500ms and 750ms after a key press and lasts for 3000 ms (step 2) while the fixation cross remains on top of the stimulus image. At step 3, subjects respond to the presence/absence of the target object. Image size is 652 by 434 pixels and the remaining of the monitor screen is filled with solid gray background

Each image was a picture of a desk, a monitor, a keyboard, a cup and several other distractor objects (See Fig. 1). The cup and keyboard were always flickering (on and off) with tagged frequencies of 12.16 and 14.19 Hz (counterbalanced between subjects). The cup and keyboard were selected as the frequency tagged items to include one object (keyboard) that contained contextually relevant information about the location of the target (mouse) and one object (cup) which was contextually irrelevant to the target (mouse) location.

The target object remained the same throughout the block. In blocks 1, 3 and 5, the target was a computer mouse while in blocks 2,4 and 6 the target was a stapler. The stapler blocks served as a control condition to isolate the effects of context from those of image content. Before the start of each block, the instruction on the screen announced the target object. The name of the target object was also displayed (as a reminder) in the resting period before each trial begins.

In blocks where the target object was a computer mouse, the target object was present in 48 images (50% of trials) located in its contextually relevant location close to the keyboard on the right side. In 5 other images (5.2% of trials), the mouse was present but in a random location out of its normal context, and the target was absent in the remaining 43 images (44.79% of trials). In other blocks where a stapler was the target object of the search, the target object was present in 48 images (50% of trials) and absent in the other 50% of trials.

### Eye tracking

In order to ensure the EEG data were not contaminated by ocular artifacts, eye position was monitored (EyeLink 1000, SR Research, sampling rate 2000 Hz). A trial was interrupted and discarded if the eye tracker detected a fixation 1.5 degrees away from the fixation cross at any time during the image presentation time. Trials were also discarded if subjects responded while images were still on the screen. At the end of each block, discarded trials were presented again, and the process repeated until all the 96 trials in a block were presented successfully.

### Behavioral measures

We analyzed performance by quantifying the fraction of target present trials for which the observer correctly decided that the target was present (hit rate) and the fraction of target absent trials for which the observer incorrectly decided that the target was present (false alarm rate). Percentage correct of trials for all trials was calculated as 0.5 hit rate + 0.5 correct rejection rate. An index of detectability (d’) was calculated from the hit rate and false alarm rate: d’ = z(HR) – z(FA) where z(.) is the z transform. Separate d’s were obtained for in-context and out-of-context conditions by using the hit rate for each condition and the same false alarm. We also measured reaction times for all trial types.

### EEG methods

Each subject’s electroencephalogram (EEG) was continuously recorded (BioSemi, Amsterdam, The Netherlands) using 32 Ag/AgCl electrodes mounted on an elastic cap and placed according to the International 10/20 System. EEG data were sampled at 512 Hz and re-referenced offline relative to the average of mastoid electrodes. Signals were band-passed filtered (between 2 and 30 Hz). Four seconds epochs of EEG data from 1 second before to 3 seconds after the stimulus onset were extracted and the baseline was removed using 100 ms before the onset of each trial. Trials that contain large amplitude artifacts (more than 80 mV absolute amplitude) at Fp1 or Fp2 channels during the presentation time, were marked and excluded.

### Time-frequency analysis and SSVEP based feature extraction

For each trial, we estimated the power spectral density in each channel at tagged frequencies (14.19 Hz 12.16 Hz) using the spectrogram of EEG data based on short-time Fourier transform as follows. Let *x*[*n*], *n* = 0,1,…, *N* − 1 be the N sample digitized EEG signal (sampling frequency = 512 Hz) during an epoch. Each epoch starts from 1.3 seconds before the stimulus onset ending at 3.29 seconds after the stimulus onset. We used a Hamming window *w*(*k*), of length L=256 data points with 50% overlap resulting in 17 overlapping sub-signals 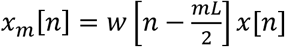. Therefore, the power of each sub-signal *x*_*m*_[*n*] at each specific frequency *f*\* can be calculated as:

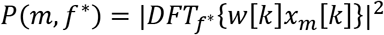

where *w*(*k*) is the Hamming window defined as 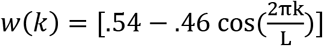 and DFT has been calculated at specific frequency *f*\* using Goertzel algorithm. Subsequently, for any given time window t_1_ to t_2_, the average power was calculated by averaging *P*(*m*, *f*\*) for all the overlapping sub-signals that fit within t_1_ to t_2_.

### Pattern Classifier

We trained and tested a binary classifier (Linear Discriminant Analysis) to classify each trial as either target present (Class 1: *signal* trials) or target absent (Class 2: *noise* trials) based on EEG spectrogram data. We chose fourteen electrodes in occipital, parietal and central parietal areas (O1, Oz, O2, PO3, PO4, P3, Pz, P4, P7, P8, CP1, CP2, CP5, CP6) a priori and based on existing knowledge that visual areas are the primary cortical source of SSVEPs (Vialatte 2010; Ding 2006). A separate classifier was designed for each time interval [*t*_1_ *t*_2_] after stimulus onset. We used the following time intervals [−1 0], [−.5 .5], [0 1], [0.5 1.5], [12], [1.5 2.5], [2 3] and [0 3] seconds for classification to investigate how the classification accuracy changes over time (with reference to stimulus onset). For each classifier, we constructed a 28 dimensional feature vector 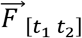 using the average power during the given time interval at two tagged frequencies of 12.16 and 14.19 Hz. LDA finds a vector of linear coefficients 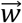 such that the linear transformation 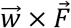, best separates the two classes. Fischer criterion is used as a measure of separability by maximizing the difference between class means normalized by a measure of the within-class scatter matrix described by the optimization function 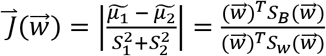, where *S*_1_ and *S*_2_ represent scatter matrices of class 1 and class 2 after projection, 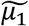 and 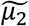 are mean of feature vectors in class 1 and class 2 after projection and *S*_*w*_ = *S*_1_ + *S*_2_ is within-class scatter matrix. We used a leave-one-out (LOO) cross-validation method to train the classifier using all but one trial and testing on the given trial. For each individual observer, we used the area under ROC curve (AUC) as a measure of classification accuracy. AUC is compared with the chance level (0.5) where higher values of AUC correspond to better classification results.

### Classifier performance evaluation

We computed the classifier performance by computing area under ROC curve (AUC) in detecting target trials for each subject and used a one sample t-test to compare group average AUCs with the chance level. In order to compare two different conditions (e.g. in-context vs. out-of-context trials), we used two sample t-test to detect significant differences between the conditions.

## Results

### Human observer performance results

Hit rate, false alarm rate, percent correct response and *d’* values for each participant were computed for each task and the applicable subset of trials in each task. Table 1 shows the average results across all 13 subjects. On average, there was no significant difference in reaction time for target mouse in-context and target mouse out-of-context trials (paired t-test, t(12)=0.95, p=0.36). However, consistent with previous studies, there was a significant decrease in hit rate for out-of-context trials relative to the in-context trials (on average 0.59 reduction in hit rate, paired t-test, t(12)=15, p=3.8 × 10^−9^). Table 1 also shows performance for the control condition in which observers searched for the presence of the stapler.

**Table 1.**
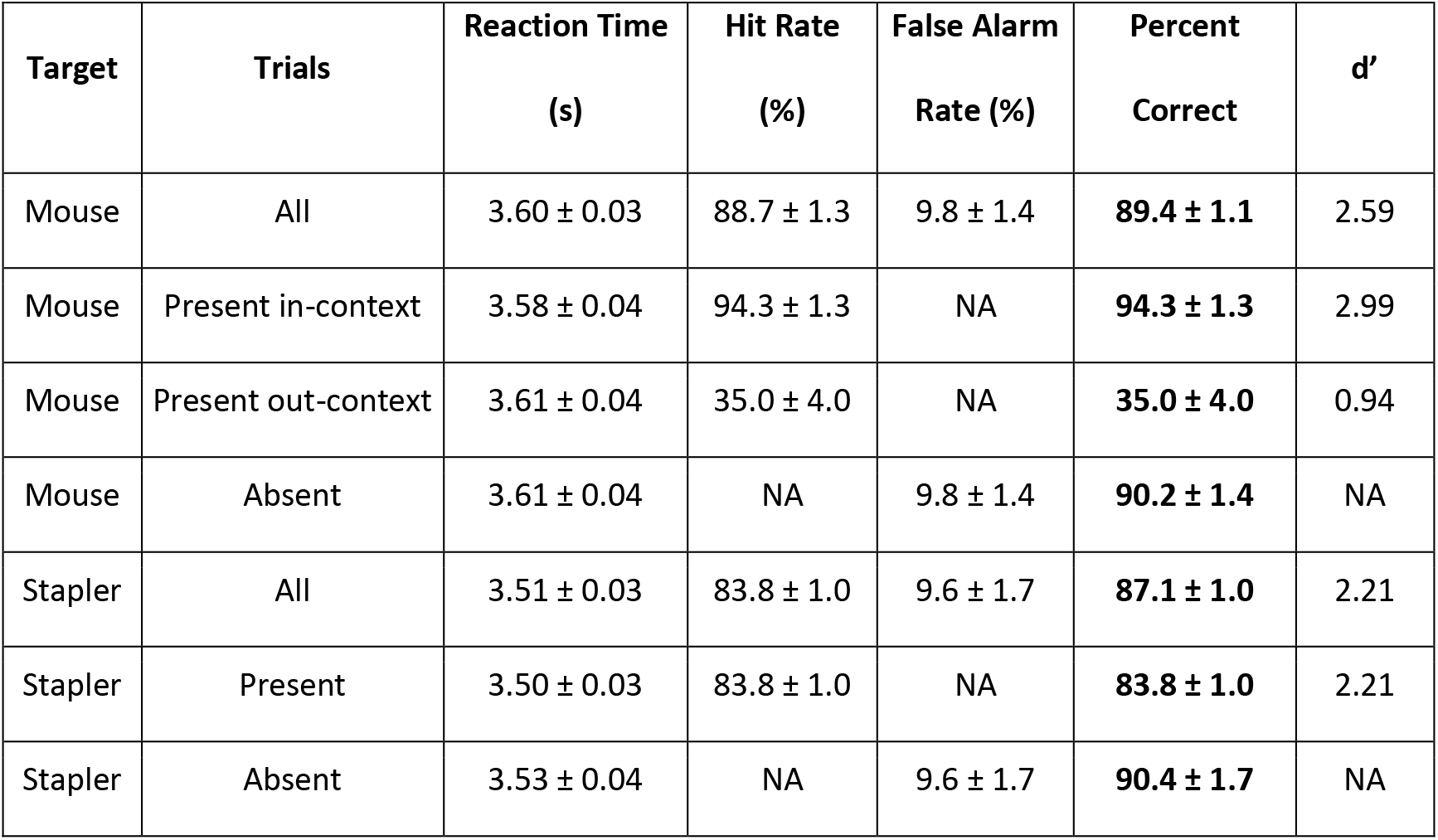
Average human observer behavioral performance.

### EEG power and scalp distribution at the tagged frequencies

EEG power at each tagged frequency and for each time window (see methods section) was computed across scalp electrodes. Due to between-subject variability in baseline values of SSVEP power, for each participant, we normalized EEG power at each electrode with respect to the total power of all 32 electrodes at the same frequency.

**Fig. 3** shows topographical maps of the average normalized power at each of the tagged frequencies during each time window averaged across all participants. The SSVEP is often maximal at occipital electrodes. However, depending on the nature of the stimulus and attention conditions, robust frequency tagged signals can also be observed at parietal-occipital and temporal electrodes (for a review see Norcia et al. 2015). Consistent with this previous work, here the largest responses at the tagged frequencies were observed at occipital, parietal-occipital, and parietal electrodes which is consistent with expected maximal locations of SSVEP and our pre-selected 14 channel locations listed in the methods section. SSVEP power increased for later the time-windows relative to stimulus onset. For example, SSVEP power was higher at the [2,3 s] time window when compared to the [0,1 s] time window.

**Fig. 3.**
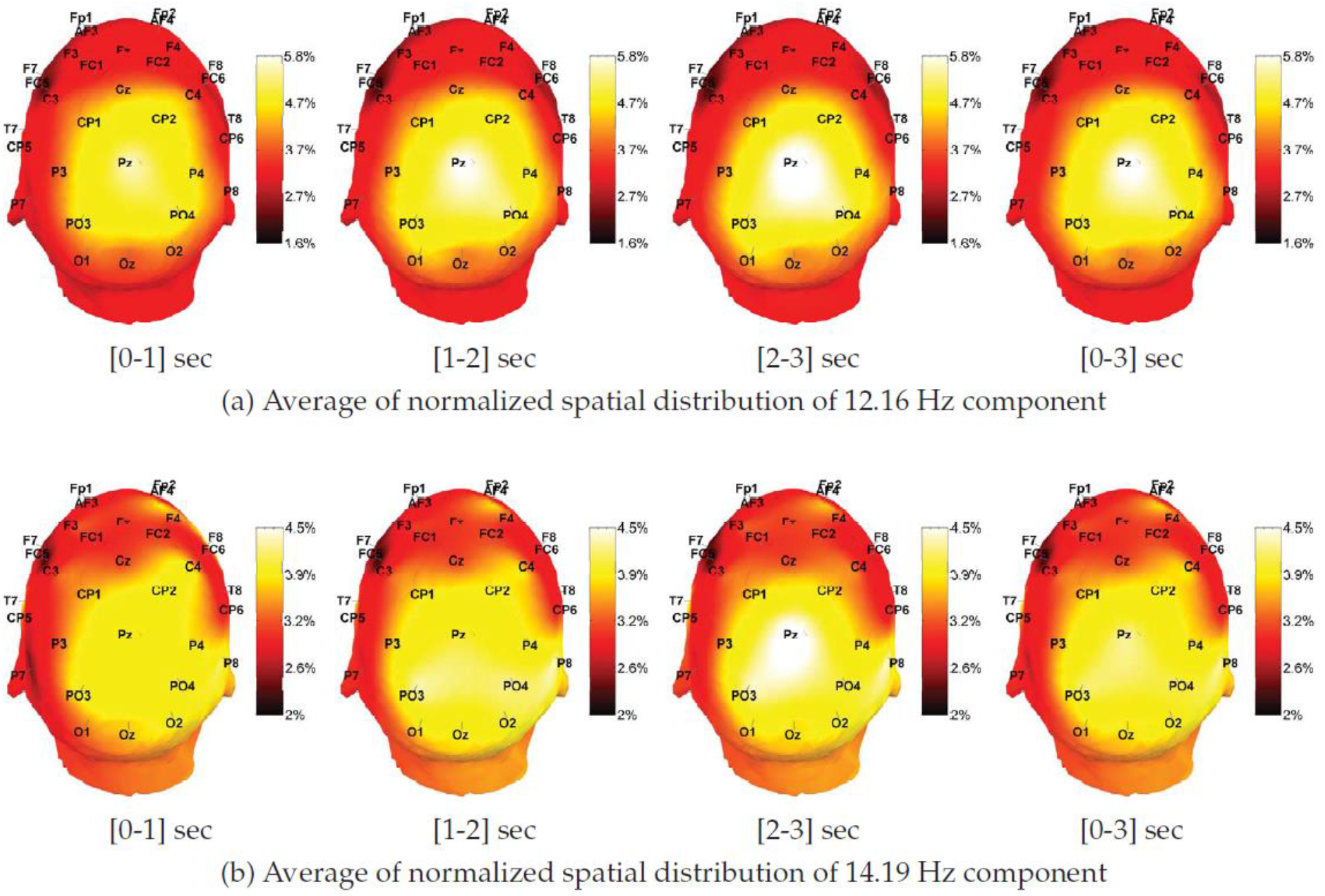
Topographical maps representing the average distribution of EEG power at both tagged frequencies (12.16 Hz and 14.19 Hz) at each of the 32 channels. The value at each channel represents the percentage of total power at the specified channel. A uniform distribution would result in 3.13% value at all locations

Fig. **4** shows a heat map of the normalized power at each channel for each subject with pre-selected channels show in the middle.

**Fig. 4.**
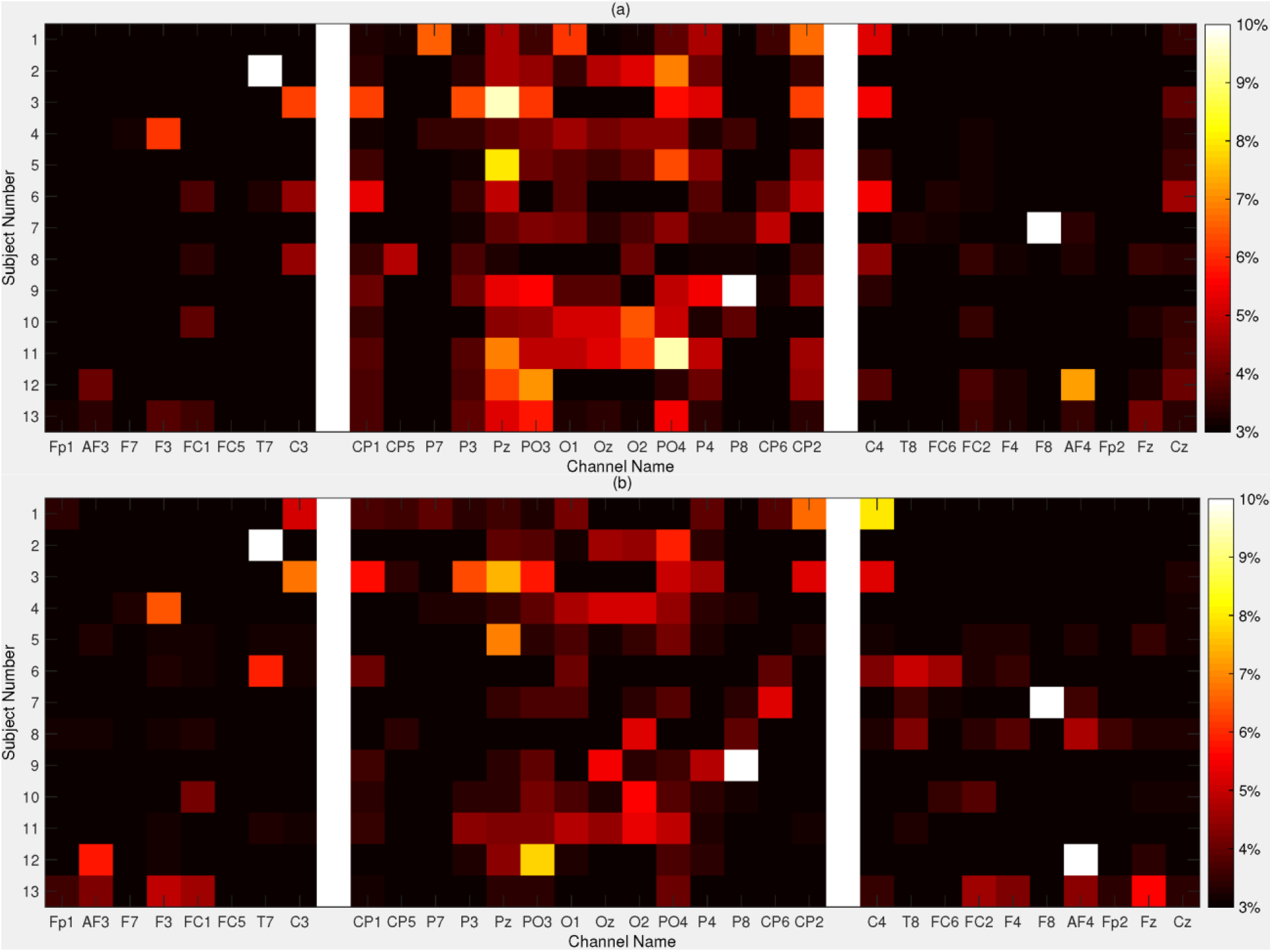
Spatial distribution of the SSVEP power for each subject plotted for (a) 12.16 Hz and (b) 14.19 Hz components. Heat map visualization shows the percentage of SSVEP power at each channel location. Values smaller than 3% and larger than 10% are colored as black and white, respectively. The middle band shows the channels that have been used for classification

### Classification performance

Classifier AUC for discriminating between target present and target absent trials using SSVEP power at the time window [0-3] post-stimulus is shown in Figure 5 for each observer and averaged across the observers. Average AUC across observers was significantly above chance level (*t(12)*=*4.37, p*<*0.001*).

**Fig. 5.**
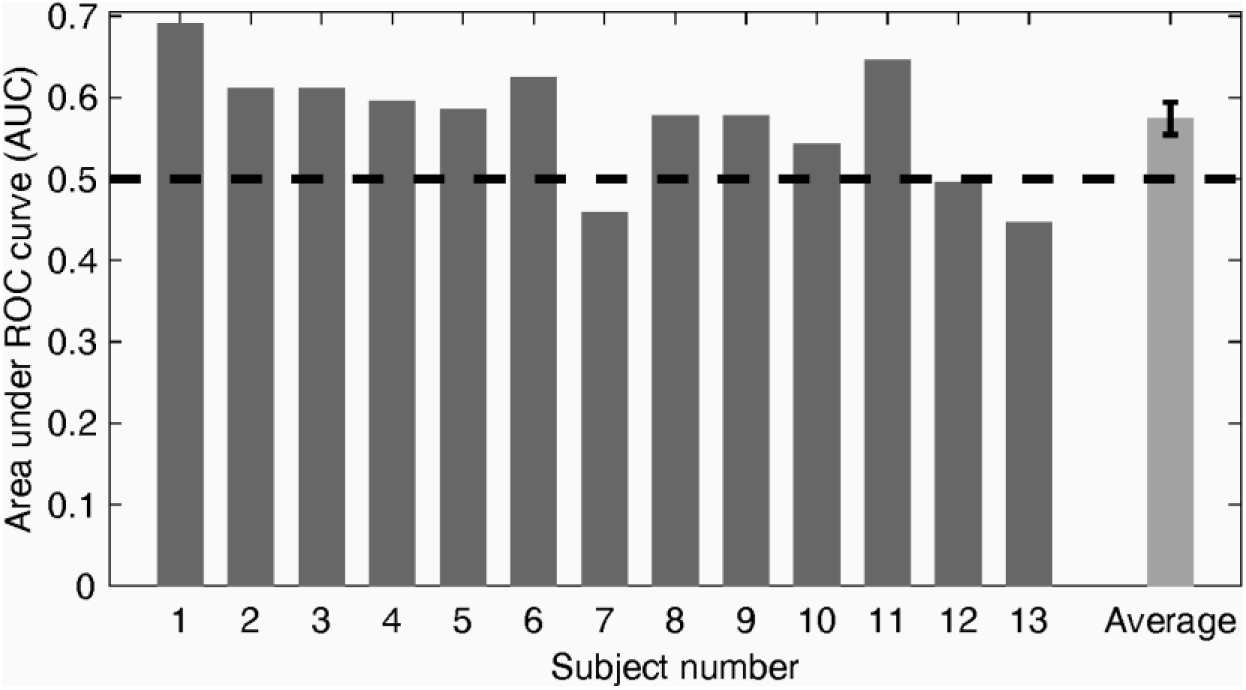
Classifier performance (AUC) detecting the presence of the target from each participant’s EEG. mean = 0.583, p<0.001, t(12)=4.37, (error bars mark +/− SEM)

In order to demonstrate the effect of stimulus presentation on classification accuracy, the same classification approach was repeated, except using EEG data in one-second-long overlapping time windows. Fig. **6** shows the average classifier performance across all subjects, plotted as a function of time-window. As expected for time windows at and before stimulus onset, AUC did not differ from chance. However, for time windows 1.5 seconds after the stimulus onset, AUC was significantly greater than chance (t(12)=2.98, p=.0115; t(12)=5.00, p=.0003; and t(12)=4.21, p=.0012) at time windows [1-2], [1.5-2.5] and [2-3] seconds post-stimulus, respectively.

**Fig. 6.**
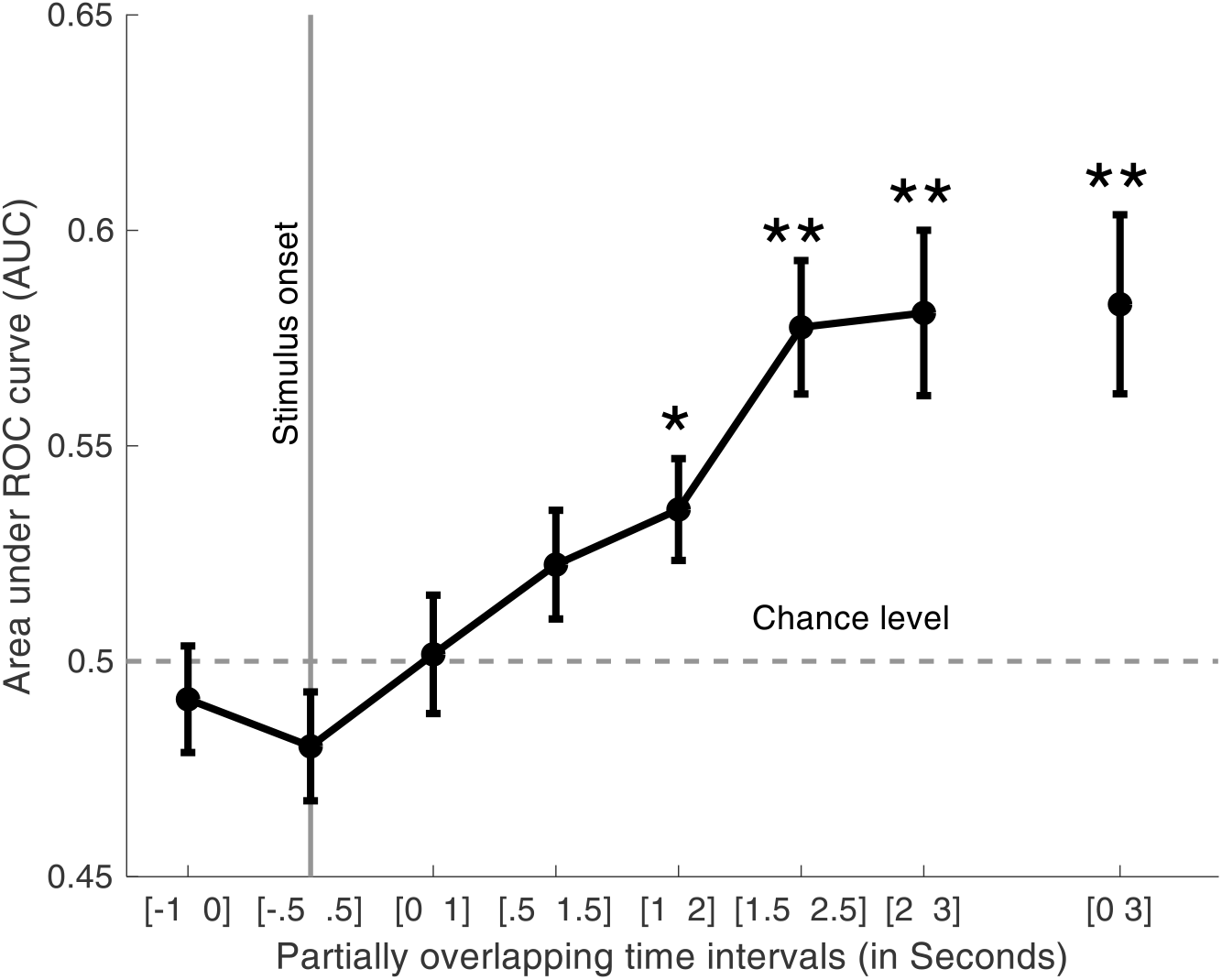
Average classifier performance detecting the presence of the target object as a function of partially overlapping time intervals from stimulus onset used for classification. (error bars mark +/− SEM). AUC is significantly greater than chance *(p<.05), **(p<.01)

### Classifier performance in detecting the target based on contextual information

In order to study the role of contextual information, the classifier was tested using a) trials with target In-context (computer mouse present in its expected location close to the keyboard on the right) and b) trials with target out-of-context (computer mouse present in image but in an unexpected location, such as the other corner of the table, or behind the monitor). Fig. **7** shows the average AUC as a function of time window was significantly higher than chance level for in-context trials, while it remained at or below chance level for out-of-context trials.

**Fig. 7.**
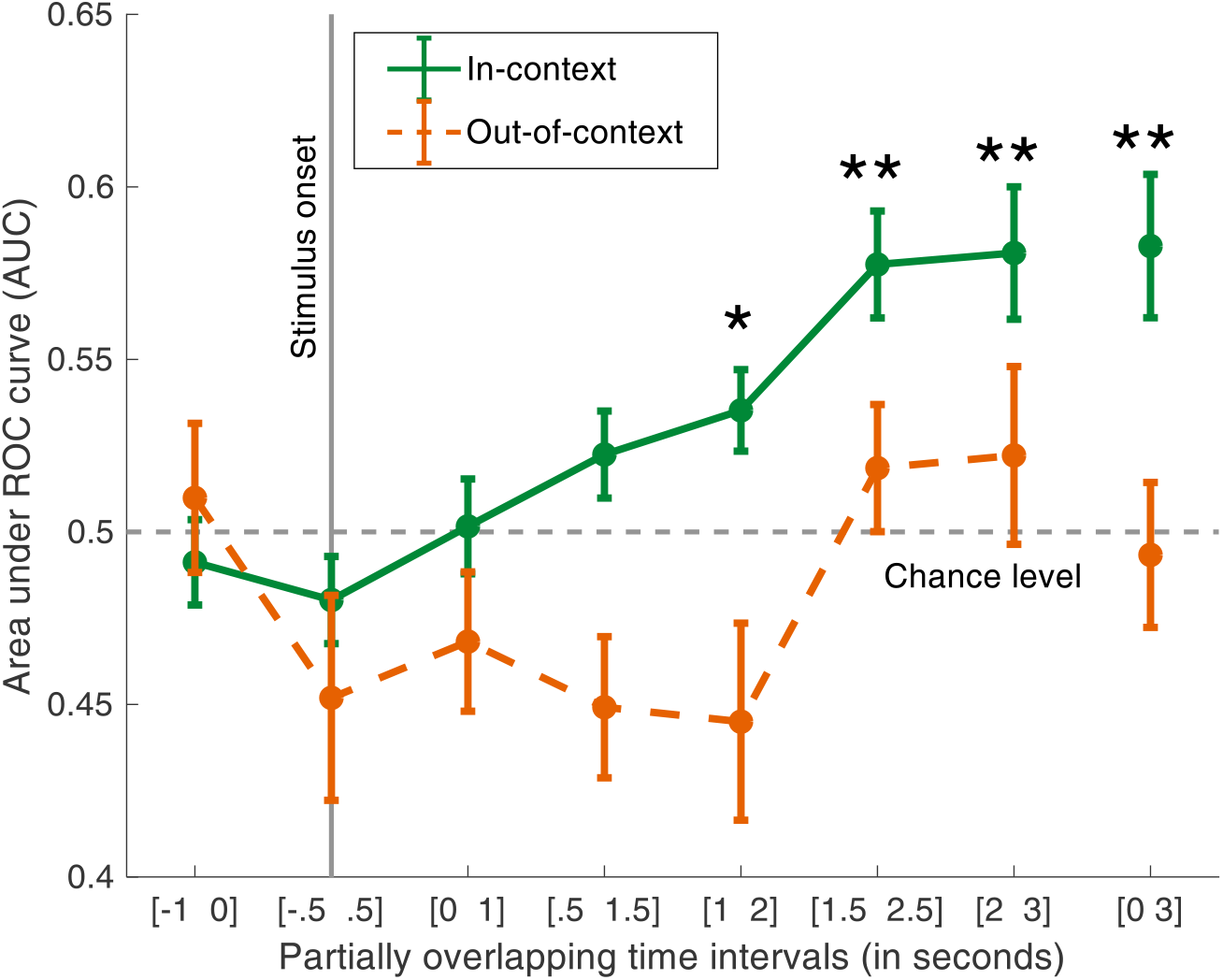
Average classifier performance for trials when the target is spatially in-context vs out-of-context. Area under ROC curve (AUC) plotted for trials with target in-context and out-of-context as a function of time intervals from stimulus onset (error bars mark +/−SEM). AUC is significantly greater than chance for target-in-context trials *(p<.05), **(p<.01)

The average classifier hit rate and false alarm rate were calculated across subjects and compared that to the average observers’ behavioral performance. Fig. **8** shows the average hit rate and false alarm rates plotted for observers’ behavior and pattern classifiers based on the SSVEPs. The observers’ performance was (as expected) higher than the classifier. However, in both cases, the difference between hit rate and false alarm rate was significantly higher (paired t-test) for the in-context trials compared to out-of-context trials (t(12)=2.9, p=0.014 for classifier and t(12) = 15.03, p=3.8×10^−9^ for observers’ performance). Overall there was no significant correlation between observers’ hit rate and classifier hit rate either for out-of-context trials (r(12)=−0.38, p=0.19) or for in-context trials (r(12)=−0.39, p=0.18). The correlation between observers’ and classifier percent correct response did not reach a significant level (r= 0.51, p=0.078).

**Fig. 8.**
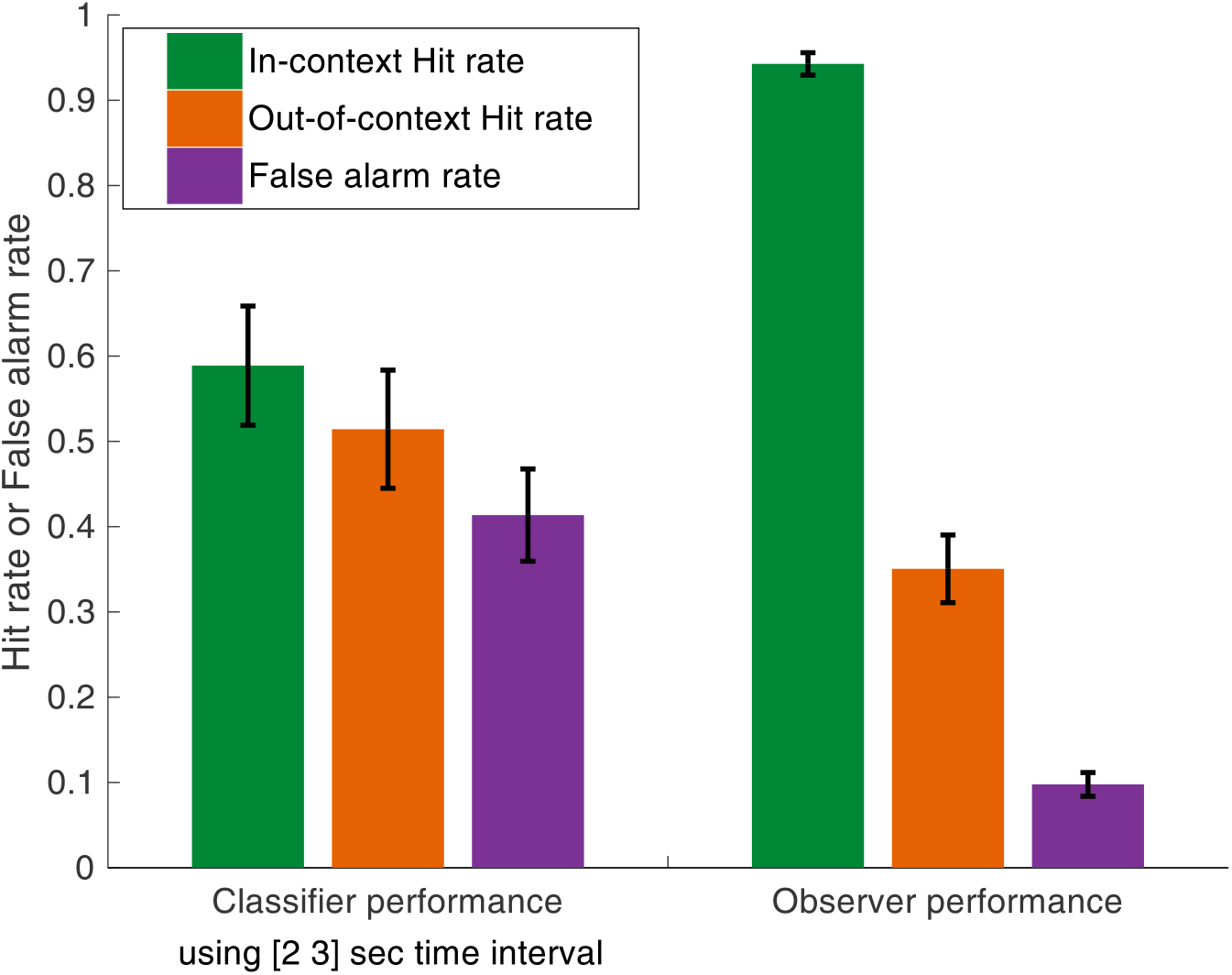
Average Hit rate and False alarm rate detecting the target for the classifier based on observer’s EEG (left) and observer behavioral decisions (right). Error bars mark +/−SEM

### Effect of target and other flickering objects’ retinal eccentricity

We investigated the effect of eccentricity on the classifier performance by computing the average hit rate corresponding to each target-present image as a function of its retinal eccentricity (Fig. **9**). For each image, target eccentricity is the distance between the target and the fixation cross, measured in units of degrees of visual angle. Across all target-present stimulus images, there was small (r=0.29) but significant correlation between target eccentricity and the average classifier hit rate, (r=0.29, p=0.038, df=51).

**Fig. 9.**
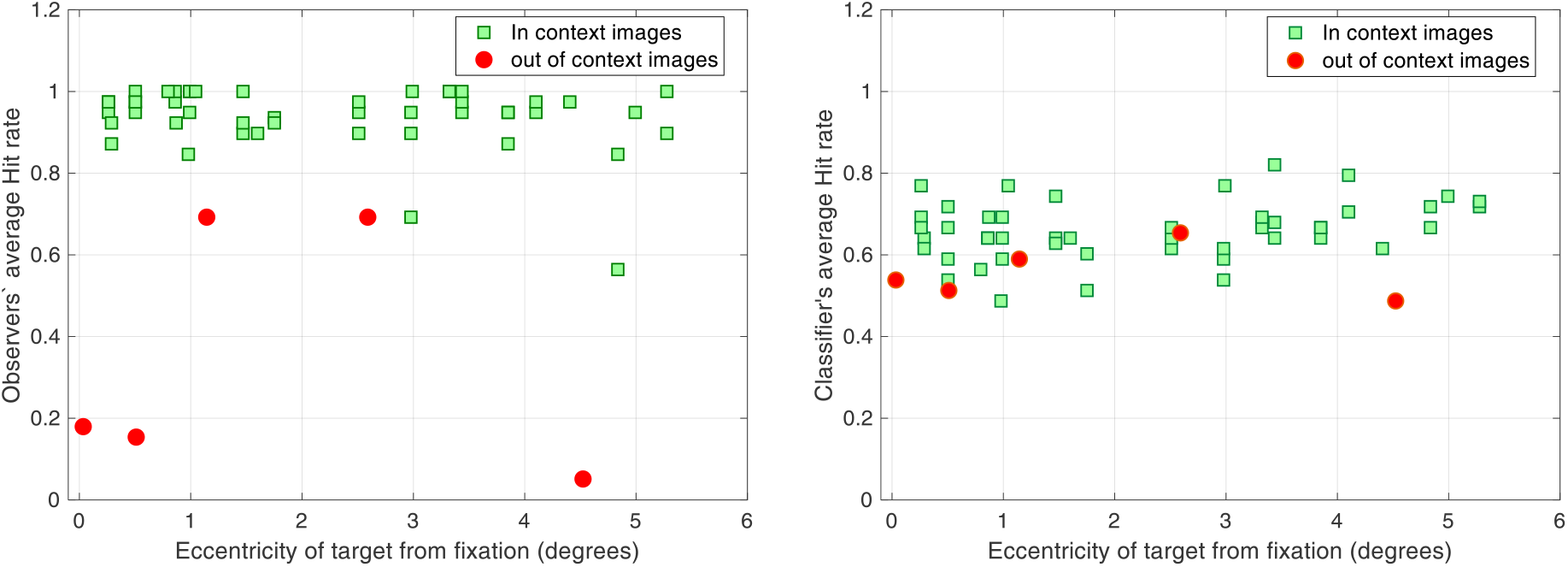
Hit rate vs retinal eccentricity of the target. Average Hit rate across observers (left figure) and classifier (right figure) for each image vs the eccentricity of the target object from the fixation

### Possible residual stimulus confounds

Although we controlled for many variables related to the target (salience with respect to the background, average retinal eccentricity) across the in-context and out of context conditions, there is always a possibility that some other property differed across the two image sets influenced the pattern classifier results. To assess any potential residual confounds, we analyzed classifier target detection accuracy for the control condition that utilized the same images as in the mouse search but for which observers searched for a stapler. If the mouse in-context vs. out of context influences on EEG responses are due to some confounding property related to the physical differences in the stimuli and orthogonal to the contextual location of the mouse then classifier detection accuracy for the stapler should also be modulated across the mouse in-context vs. out of context images. However, there was no difference in stapler classifier detection as a function of the contextual location of the mouse (see Fig. **10**). The overall AUC for detecting stapler did not significantly differ between mouse-in-context and mouse-out-of-context trials (t(24)=−1.4, p=0.17), between mouse in-context and mouse absent trials (t(24)=0.56, p=0.58) or between mouse out-of-context and mouse absent trials (t(24)=1.73, p=0.09). These results provide further evidence that the classifier target accuracy for the mouse is related to its contextual location and not any uncontrolled image-specific physical property.

**Fig. 10.**
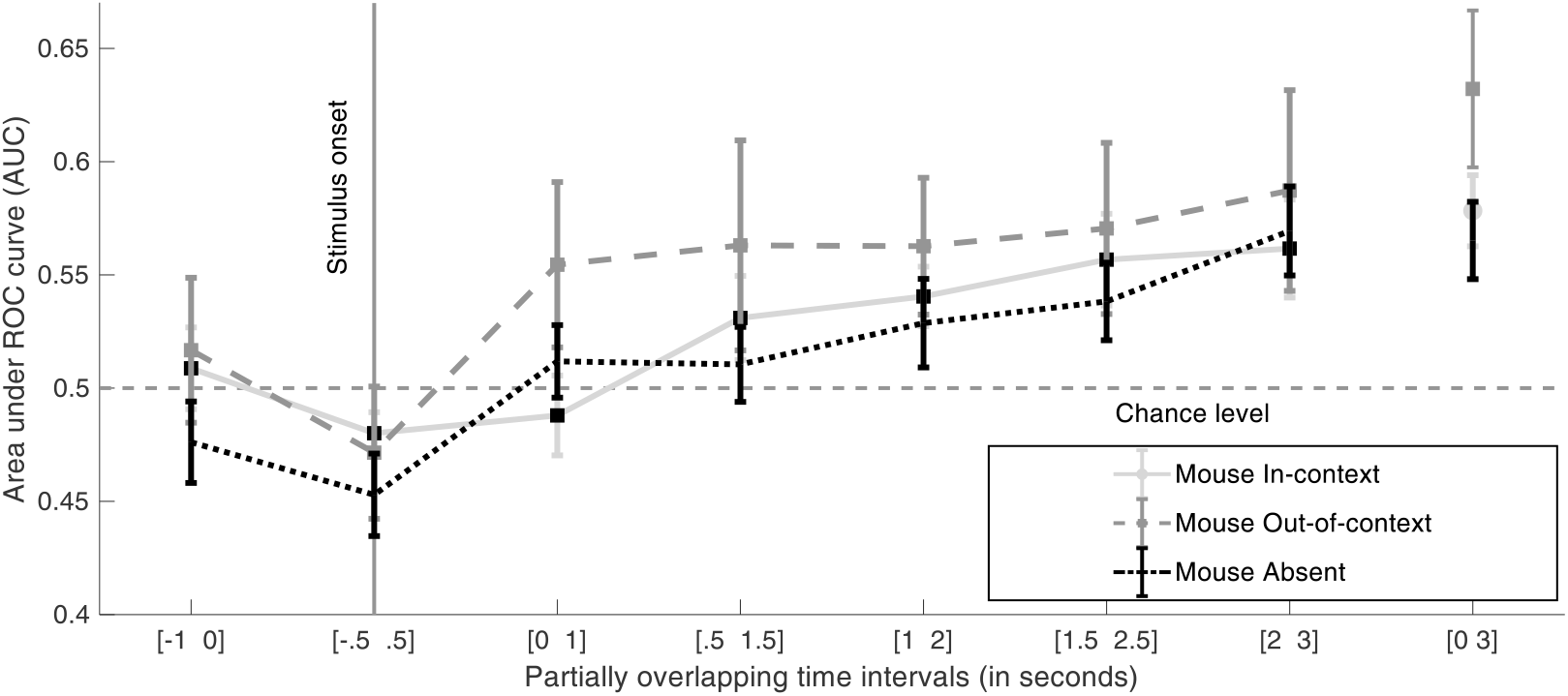
Average classifier performance in detecting stapler applied the same set of images as those utilized for the mouse detection trials. Area under ROC curve (AUC) plotted for trials with mouse In-context, Out-of-context and absent as a function of partially overlapping time intervals from stimulus onset

### Comparison between EEG classifier and observers’ aggregate responses

To evaluate whether there is a relationship between observers’ behavior and brain activity, we compared the performance of the classifier and an aggregate of observers’ behavior for each individual stimulus image. Fig. **11** shows the proportion of trials classified as target present by the classifier as a function of observer decisions. The top right corner (and the bottom left corner) of the scatter plot show stimulus images correctly classified by both the classifier and observers as target present (Hits) and absent (correct rejections). The horizontal dotted line represents the chance level of 55% hit rate as 55% of images are target present (50% In-context and 5% Out-of-context). Both the classifier and the observer performed poorly on target out-of-context images.

**Fig. 11.**
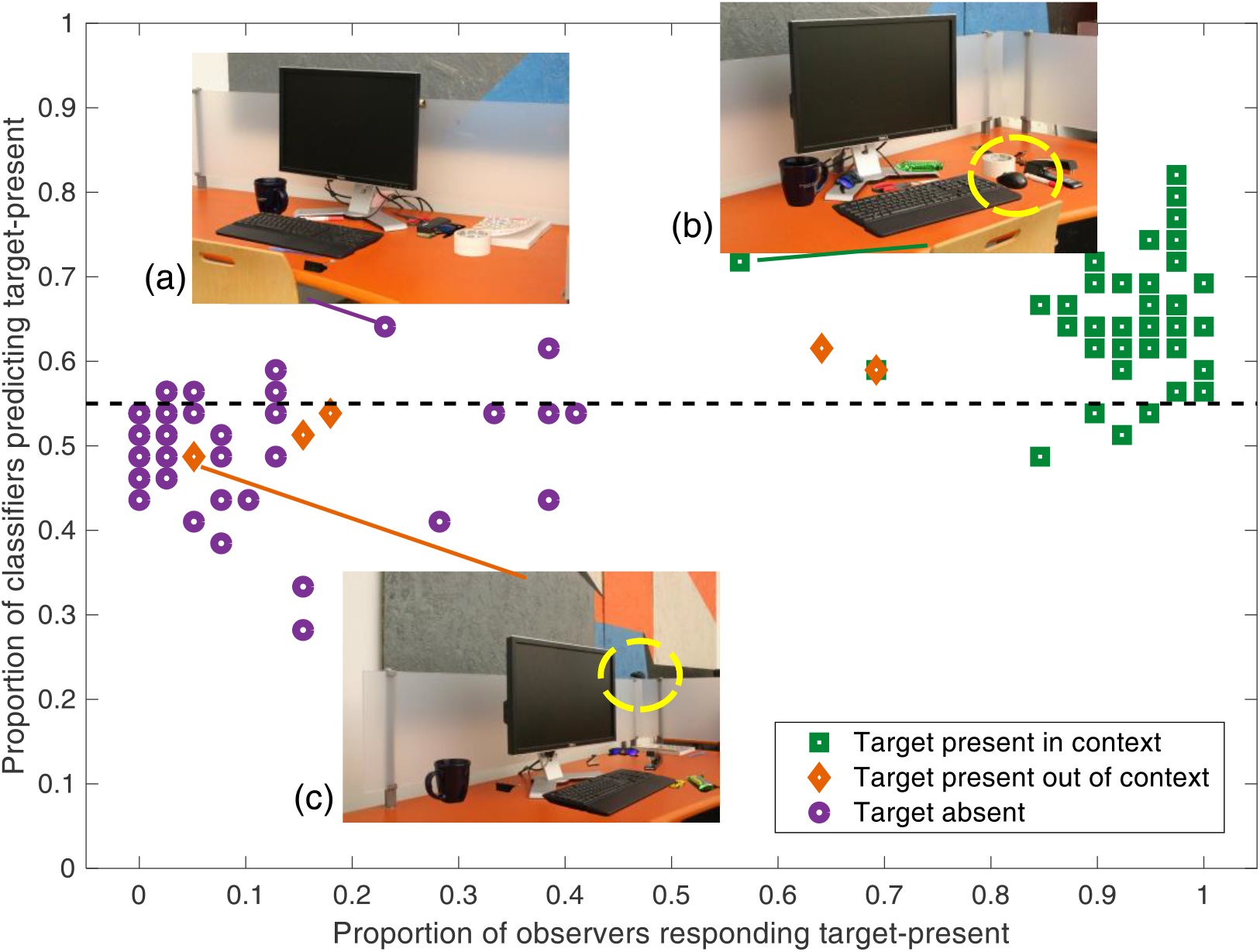
Relating classifier performance to observer behavioral performance: Each data point represents a stimulus image. The scatter plot shows the proportion of trials for which the classifier predicts “target present” vs proportion of trials for which observers responded “target present”. The horizontal dotted line shows classifier chance performance corresponding to prior probability (55%) of target present detection (50% in-context and 5% out-of-context). Sample images with target absent (a), target present In-context (b) and target Out-of-context (c) have been marked and shown. Yellow dotted circles in images (b) and (c) show the location of the target and are not part of the stimulus image

## Discussion

Contextual relationships among objects in scenes are considered to be a fundamental property used by the human brain to guide visual search (Eckstein 2017; Wolfe and Horwitz 2017; Wolfe et al. 2011, Võ et al. 2019). The majority of the EEG studies have concentrated on assessing how neural signals are affected by the consistency or inconsistency of an object with the background scene (Martens et al. 2011). Demiral, Malcolm, and Henderson (2012) varied the spatial congruency of objects in scenes and used EEG to measure the semantic mismatch event-related potential (ERP) known as the N400 (Kutas and Hillyard 1980) with a 300ms scene preview. Spatially incongruent objects led to a robust N400-like modulation that was weakened when object and scene were presented simultaneously. Vo and Wolfe (2013) found a clear dissociation between two types of inconsistencies in scenes. Semantic scene inconsistencies refer to objects that do not typically belong to the scene (e.g., a motorcycle in a bedroom). Semantic scene inconsistencies produced negative deflections in the ERP in the typical N400 time window. Syntactic scene inconsistencies refer to objects that typically appear in the scene but that are placed at an unlikely location (e.g., slippers on the bed). Vo and Wolfe found that mild syntactic scene inconsistencies elicited a late positivity resembling the P600 that is typically found for syntactic inconsistencies in sentence processing. Extreme syntactic violations (e.g., a hovering beer bottle defying gravity) were associated with earlier perceptual processing difficulties reflected in a negative deflection in the N300-N400 time-window but failed to produce a P600 effect.

These studies are critical in understanding the neural signatures of object/scene consistency but do not isolate how target-related neural activity is modulated by the contextual location of the target in the scene and how the activity relates to search decisions. Previous studies have demonstrated the influence of context on both target-related activity EEG and fMRI signals for synthetic displays (Johnson et al. 2007; Giesbrecht et al. 2013; Kasper et al 2015; Greene et al 2007). A previous study utilizing fMRI (Preston et al. 2013) decoded target-related activity during search with real scenes in the intra-parietal sulcus (IPS) and the frontal eye fields (FEF) but did not evaluate the modulation by scene context. Another study showed using decoding methods that the coarse expected location of a target is represented in the lateral occipital complex (LOC) and IPS (Guo et al. 2012). A recent study (Brandman and Peelen 2017) utilized MEG and fMRI to show that decoding of object categorization (animate vs. inanimate) in the lateral occipital area and posterior fusiform sulcus (pFs) improved when a degraded image of an object was presented within a consistent scene. Unlike this latter study, the current study involved visual search for an object that might be present or absent in the scene. We specifically investigated the influence of placing the object in unexpected spatial locations. We were particularly interested in the influence of contextual location on the accuracy of a target detection classifier based on the SSVEPs.

We found that the contextual location of the searched target influenced pattern classifier performance detecting the target using frequency tagged evoked brain responses. The accuracy decoding the presence of the search target decreased when the object was placed out of context. Attributing the modulation of EEG activity to the contextual location of the target requires careful control of possible confounding variables arising including target saliency, retinal eccentricity, and eye movements. The dissociation in classifier accuracy across contextual location cannot be attributed to a variety of factors we controlled for. We matched the average retinal eccentricity of the target across images for the in-context and out-of-context conditions. Our design controlled for eye movements utilizing a gaze-contingent design to control for possible contamination of the neural signals from oculomotor commands. The retinal eccentricities of the temporally modulated objects (non-targets) providing the EEG signals were also matched across conditions by varying the fixation cross across trials. Target saliency against the surrounding background and/or other image properties that might have differed across contextual conditions cannot be used to explain our results. By design, we placed the target with a similar surrounding local background (orange desk). Critically, if the dissociation in decoding accuracy of the presence of the target object (mouse) across context conditions was related to some low-level physical differences across images, then we should expect that to show an accuracy dissociation even when observers view the same images but are performing an orthogonal task. Yet, when observers searched for an unrelated object (stapler) while viewing the same images, the classifier did not find any dissociation in accuracy across in context and out of context target conditions.

We also found a relationship between the propensity of a scene to lead to target present responses for the SSVEP classifiers and that of observers’ behavioral responses. Images for which observers likely detected the target were also more likely to lead to a correct detection for the pattern classifiers. Thus, the results suggest that the identified target-related neural activity is modulated by the variations in task difficulty from image to image in a similar manner as the behavioral observer decisions and is consistent with previous studies showing a relationship between the image-specific behavior and neural signals in EEG and fMRI (Das, Giesbrecht, & Eckstein, 2010; Guo et al., 2012).

There was a small but significant effect of target eccentricity across images on decoding accuracy for the eccentricity ranges in our study (Regan, 1966; Ding et al., 2006; Lin et al., 2012; Meredith and Celesia, 1982). The smaller effects relative to previous studies (e.g., Ding et al., 2006; Lin et al., 2012) might be related to the lower target eccentricities used here (less than six degrees of visual filed).

To summarize, our study finds consistent evidence that target-related EEG activity is modulated by scene context during target search with real world scenes. The results add to a growing literature showing how spatial relationships between an object and scene alter neural activity that represents object/scene inconsistencies and that codes the presence of the searched target. The proposed paradigm might be used for future studies attempting to partition different components of contextual information such as the consistency with the background, the co-occurring object most predictive of the target location and the spatial configuration of other objects in the scene.

## Acknowledgments

The first author is currently with Advanced Brain Monitoring Inc, Carlsbad, CA, 92008

## Declarations

### Ethics approval

The experimental protocol was approved by the institution review board (UCSB Human Subjects Committee, PROTOCOL NUMBER 15-18-0468).

### Consent to participate

All participants signed a consent form approved by the IRB.

### Consent for publication

All participants signed a consent form approved by the IRB.

### Availability of data and material

Raw data and materials are provided at the following link:

*Meghdadi, Amir (2020), “EEG signatures of contextual influences on*
*visual search with real scenes”, Mendeley Data, v3*
*http://dx.doi.org/10.17632/4t6hjv3cm6.3*

### Code availability

Not currently available. Will be provided upon request.

### Authors’ contributions

A.M. implemented the tasks, conducted the experiments, collected the data and performed processing and analysis of the data. M.E. conceived the original idea, oversaw the experimental protocols and data analysis and contributed to the design of the approaches to the analysis. B.G. oversaw data collection and contributed to the design of the experimental protocols. A.M. M.E. and B.G. wrote the manuscript.

